# Mice use predictive environmental cues to adapt predatory behavior in hunting tasks of variable difficulty

**DOI:** 10.64898/2026.09.03.749200

**Authors:** Jacob L. Amme, John M. Grady, Kiran Bhaskaran-Nair, Varun Sinha, Samuel J. Brunwasser, Anthony I. Dell, Keith B. Hengen

## Abstract

Prey capture has long served as a model of complex, multimodal behavior with ethological relevance. In this context, the ability of laboratory animals to capture fleeing prey has been investigated in terms of the sensory systems and neuroanatomical regions required for successful capture. While the ability to hunt is experience-dependent, prey capture is often assumed to be a reflexive behavior, governed by innate neural circuits and a basic drive to obtain food. Whether hunting performance depends on complex learning and cognitive strategy remains largely untested, in part because it is difficult to systematically manipulate the difficulty of predator-prey interactions. To address this, we varied ambient temperature across thousands of trials in which laboratory mice pursued cold-blooded cockroaches, whose movement speed was directly determined by body temperature. As expected, capture difficulty increased as a function of temperature. To test whether mice rely on external cues to adjust their predatory behavior, we then mismatched the ambient temperature and that of the prey, finding that mice learn to use relevant environmental information to inform their hunting strategy. Our results suggest that animals can extract and apply meaningful information from predictive environmental cues to adapt their learned behavior.

## Introduction

The struggle to acquire food is one of the strongest and most fundamental drivers of natural selection^1–4^. Animals that hunt other organisms have evolved a sophisticated set of circuits and systems optimized for the identification, pursuit, and capture of prey, thereby forming a central locus of brain function^4–6^. In predatory mammals, tracking prey involves integrating multiple sensory modalities including vision, audition, olfaction, and somatosensation^4,6,7^. Moreover, to successfully pursue and catch fleeing prey, predators must employ complex systems of motor control in a goal-directed manner, while also navigating heterogeneous and often unpredictable environmental conditions^2,6^. In spite of these complexities, prey capture is generally understood to be an innate, reflexive behavior that, like courtship displays, nesting instinct, and birdsong production, arises from pre-programmed circuitry that is fine-tuned by experience^8,9^. Yet even innate behaviors require some flexibility, enabling animals to adjust their actions to the shifting requirements of real-world environments^10^. For predators, such adaptability is essential: variable conditions and prey dynamics invariably alter the functional demands of the hunt. Despite this, it is largely unknown how external environmental context shapes the expression of innate predatory hunting behavior.

An animal’s basic need to generate behaviors suitable to the context of its environment drives the evolution of the central nervous system^11–14^, making the study of natural behavior critical for understanding how the brain operates — from sensory processing to higher-order cognition and motor control^15–17^. Predatory hunting, defined as the pursuit, capture, and killing of animals for food, is highly conserved and widely expressed across the animal kingdom, making it a useful paradigm for studying complex, ethologically relevant behavior^2–4^. Mice, like most rodents, are natural hunters^18,19^ and readily learn to hunt insects in a laboratory setting^20–23^, making them an ideal model organism for studying predatory behavior and its neural correlates in controlled experimental conditions. In contrast to artificial tasks that require extensive training or engineered stimuli, mouse prey capture is rapidly acquired and engages natural sensory cues^24–28^. Mice primarily rely on vision to hunt insects in lighted conditions^24,26,29–32^ but may also utilize auditory, olfactory, and somatosensory information to help detect, track, and ultimately subdue their prey^25,27,33,34^. Recent studies have begun to reveal the neural circuitry that mediates the sensory processing, sensorimotor transformations, predatory drive, and encoding of hunting actions^25,26,34–42^. Yet, despite detailed insight into its underlying endogenous circuitry, the extent to which mouse predatory hunting behavior is modulated according to context and experience remains unclear.

As a core element of natural behavior across taxa, predatory hunting is often treated as a fixed action pattern — a closed behavioral program that simply relies on innate circuitry^11^ — yet its expression is more nuanced than this view suggests. Conventionally viewed, predatory hunting involves a series of sequential stereotyped actions, including prey search, pursuit, attack, and consumption. Across many species, prey-hunting behavior has been shown to arise innately in response to the detection of certain prey or prey-like stimuli^35–37,43–45^. These external stimuli activate a genetically determined instinctive behavioral cascade that ultimately leads to successful prey capture or escape^6,8,9,27^. In rodents, species differ markedly in the pattern and execution of their stereotyped hunting sequences, reflecting the distinct evolutionary pressures that give rise to species-typical innate predatory behavior^23,46–50^. However, these instinctive action patterns are not entirely rigid; their expression can undergo substantial refinement through experience^51^. Effective predatory hunting in mice involves extensive learning, with naive laboratory mice showing an order-of-magnitude reduction in time to capture over repeated trials until achieving stable performance^24,27,28,52,53^. These improvements likely reflect experience-dependent sensorimotor adjustments in prey detection, tracking, grasping, and biting^6,27,28^ as well as motivational changes including reduced neophobia and strengthened appetitive drive resulting from exposure to a novel food reward^24,34,39,54–57^. Notably, southern grasshopper mice learn to adjust their predatory attack behavior in response to prey-specific defenses^58,59^, indicating that previous experience can play an important role in shaping hunting behavior. More recent studies have shown that prey capture experience can alter the salience of certain visual features^60^, and that it can also induce structural plasticity in the visual cortex to enhance visual function in juvenile rodents^53^. Taken together, these studies demonstrate that predatory hunting behavior is fundamentally instinctive yet also shaped by learning, with predatory experience potentially remodeling the neural circuits that support it. And while mouse predatory hunting clearly comprises both innate and learned components, whether and how this behavior is flexibly adapted to environmental conditions relevant for hunting remains poorly understood. We hypothesized that mice can learn to adjust predatory behavior in response to behaviorally-relevant environmental context. Here, we tested whether an innate, closed-program behavior — predatory hunting — can exhibit the characteristic flexibility of a plastic, open program. Specifically, we asked whether mice form abstract, indirect associations that serve as a predictive framework for adapting their predatory behavior to environmental context.

To address this question, we took advantage of the fact that movement speed is metabolically constrained by ambient temperature in cold-blooded insects. Thus, by varying ambient temperature across thousands of mouse/insect prey capture trials, we were able to directly modulate the difficulty of the hunting task via control of an environmental variable. This design dissociates two forms of adaptive behavior: a closed-program adaptation, in which mice simply react to the moment-to-moment speed of the insect, and an open-program adaptation, in which mice learn to use ambient temperature to forecast the difficulty of upcoming trials and adjust their behavior accordingly. We first measured baseline predatory learning at room temperature, then bidirectionally shifted ambient temperature and assessed the resulting changes in hunting performance. Finally, to test whether mice with experience hunting across a wide range of temperatures might use temperature cues to adapt their hunting behavior, we mismatched the temperature of the mouse and insect, finding that hunting performance was altered when the predictive value of the environmental context was removed. Our results reveal that mice adapt innate, closed-program behavior to contextual environmental structure, deploying learned associations to anticipate task demands in a manner characteristic of open behavioral programs.

## Results

Recent studies of mouse predatory hunting have carefully identified innate neural circuitry that supports this natural behavior^25,26,34–42^. Although this work has advanced our understanding of how hunting behavior is encoded in the mouse brain, far less is known about how context and experience shape its expression. Mice reliably improve their prey capture performance with practice^24,27,28,52,53^, yet it remains unclear whether their hunting behavior exhibits broader behavioral plasticity. In particular, the extent to which mice adjust their hunting strategy and behavior under varying environmental conditions remains unknown. To investigate this question, we developed a novel mouse/insect prey capture assay in which we bidirectionally manipulated ambient temperature, allowing us to systematically vary hunting difficulty in a context-dependent manner (**Fig. 1**). In this experimental paradigm, we separately housed C57BL/6 laboratory mice (*Mus musculus*) and red runner cockroaches (*Shelfordella lateralis*) within temperature-controlled chambers (**Fig. 1**). Mice were housed individually in a home cage that also served as the hunting arena. Using overhead video cameras with markerless pose estimation^61^ we tracked and quantified mouse and red runner movement before and during trials, which were conducted at five target temperatures ranging from 14 to 35*^◦^*C (**Fig. 1a,b**). For each block of trials, we set the chamber to the target temperature and maintained it at steady state for 30 minutes before beginning the trials, ensuring sufficient time for red runner body temperature to passively adjust to the ambient conditions (**Fig. 1b**).

**Figure 1:**
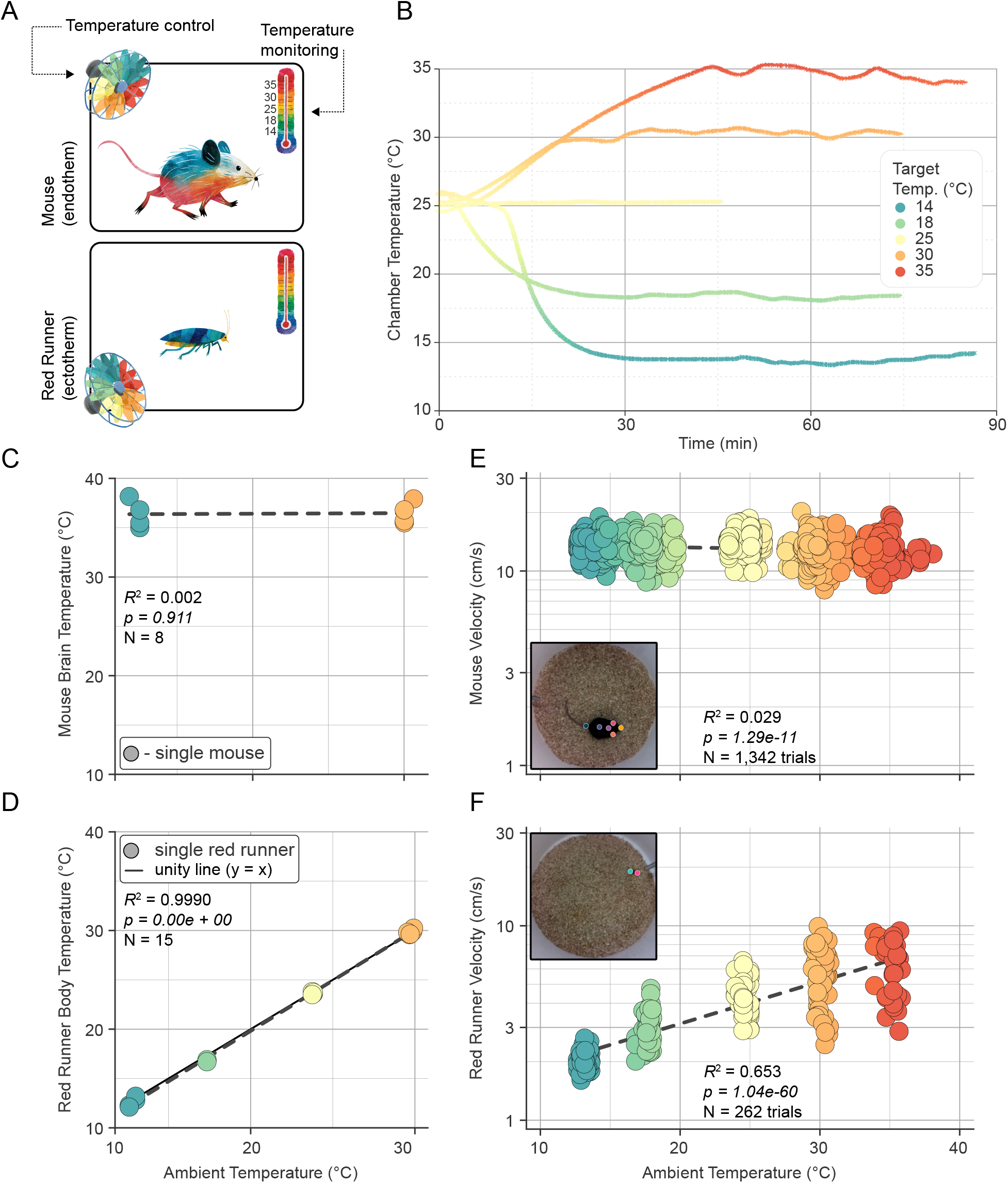
A temperature-controlled assay for bidirectionally modulating prey-capture difficulty. **(A)** Schematic of the experimental paradigm. C57BL/6 mice (*Mus musculus*, endotherm) and red runner cockroaches (*Shelfordella lateralis*, ectotherm) were separately housed in temperature-controlled chambers. Each mouse’s home cage served as the hunting arena. Chamber temperature was regulated and continuously monitored, with overhead video used for markerless pose estimation of both species. **(B)** Chamber temperature over time for the five target temperatures (14, 18, 25, 30, 35*^◦^*C). Chambers reached and maintained steady state within *∼*30 min, and were held for 30 min before trials began to allow red runner body temperature to passively equilibrate to ambient conditions. **(C)** Mouse brain temperature (*n* = 8 mice) and **(D)** red runner body temperature (*n* = 15 red runners) as a function of ambient temperature. Each data point represents a single individual animal. Dashed trendlines represent Ordinary Least Squares (OLS) linear regression fits (*body temp ∼ trial temp*). Mouse brain temperature remained invariant to ambient temperature (*y* = 0.0059*x* + 36.30, *R*^2^ = 0.002, *p* = 0.911), whereas red runner body temperature closely tracked ambient temperature (*y* = 1.02*x −* 0.61, *R*^2^ = 0.999, *p <* 0.001; solid line denotes *y* = *x*). **(E)** Solitary mouse median trial speed as a function of ambient temperature (*n* = 7 mice, 191.7 *±* 15.7 trials per mouse, *n* = 1, 342 total trials). Log-transformed speeds were analyzed using a Linear Mixed-Effects Model (LMM) with animal ID as a random intercept (*log*(*vel*) *∼ trial temp* + (1*|animal*)). Mouse locomotor speed was largely invariant to ambient temperature (marginal *R_m_*^2^ = 0.029, *p <* 0.001), after accounting for significant baseline differences between individual animals (random intercept, *p <* 0.001). Biologically, while mouse identity dictates baseline locomotor performance, ambient temperature in isolation does not systematically drive mouse speed. **(F)** Solitary red runner median speed as a function of ambient temperature (*n* = 262 total trials). Each data point represents a single individual red runner. Log-transformed speeds were evaluated using a multiple OLS regression controlling for log-transformed body mass (*log*(*vel*) *∼ trial temp* + *log*(*mass*)). Red runner locomotor speed increased steeply with temperature (*R*^2^ = 0.653, *p <* 0.001), after controlling for body mass, which was not a statistically significant predictor (*p* = 0.156; fit line shown at median mass). Biologically, thermal conditions strongly govern ectothermic movement regardless of body size. Insets show representative overhead video frames with markerless pose estimates. Point color denotes mean trial temperature throughout.

After confirming that red runner body temperature reliably tracked ambient temperature under this protocol (**Fig. 1c,d**), we then assessed solitary mouse and red runner movements at the five target temperatures between 14 and 35*^◦^*C. We predicted that, consistent with a general understanding of warm-blooded and cold-blooded animals, the movement rate of warm-blooded mice would be unaffected by ambient temperature, whereas the movement rate of cold-blooded red runners would increase with higher temperatures, at least when the species were monitored in isolation. As expected, we found that (in isolation) mouse locomotor speed and acceleration remained invariant to ambient temperature (**Fig. 1e**; **Fig. S1a**), with no meaningful changes in movement even at the extremes of the investigated temperature range (mouse solitary speed: *R*^2^ = 0.029, *p <* 0.001; mouse solitary acceleration: *R*^2^ = 0.000, *p* = 0.612). Conversely, red runner locomotor speed and acceleration (in isolation) was strongly dependent on temperature (**Fig. 1f**; **Fig. S1b**), with the rate of movement increasing *∼*exponentially until reaching a peak at 30*^◦^*C (red runner solitary speed: *R*^2^ = 0.653, *p <* 0.001; red runner solitary acceleration: *R*^2^ = 0.722, *p <* 0.001). Red runner speed remained stable at 35 relative to 30*^◦^*C, suggestive of a biophysical limit to the thermal sensitivity of insect metabolism. Importantly, these data suggest that there is a viable range of temperatures in which red runner speed is predictably modifiable without compromising the functionality of either species to thermal effects. Thus, using this assay, we could effectively modulate red runner movement by controlling ambient temperature, positioning us to test whether this temperature-dependent variation in prey speed influences hunting outcomes and the expression of mouse hunting behavior.

Prior to conducting hunting trials at different temperatures, we first trained naive mice to hunt at room temperature (**Fig. 2a**, top). After habituating the mice to live red runner prey (see **Methods**), we carried out 10 days of training trials at room temperature. Each trial day consisted of overnight food restriction followed by 6 consecutive red runner trials (**Fig. 2a**, bottom). Mice rapidly learned to hunt during this initial training phase, with average capture times decreasing from 508 to 26 seconds and stabilizing by the third to sixth day (**Fig. 2b**; **Fig. S2**). Using machine vision, we continuously tracked mouse and red runner movements to quantify both fine-scale kinematic features and overall trial outcomes (**Fig. 2c**). For an experienced mouse, once it detected the prey, the remainder of the trial could be segmented into discrete pursuit bouts punctuated by contacts, during which the mouse grappled with the red runner and attempted to subdue it (**Fig. 2d**). Following one or more contacts, the trial typically ended with the mouse incapacitating and consuming the red runner. These data demonstrate that mice rapidly refine a set of conserved hunting behaviors in an experience-dependent fashion.

**Figure 2:**
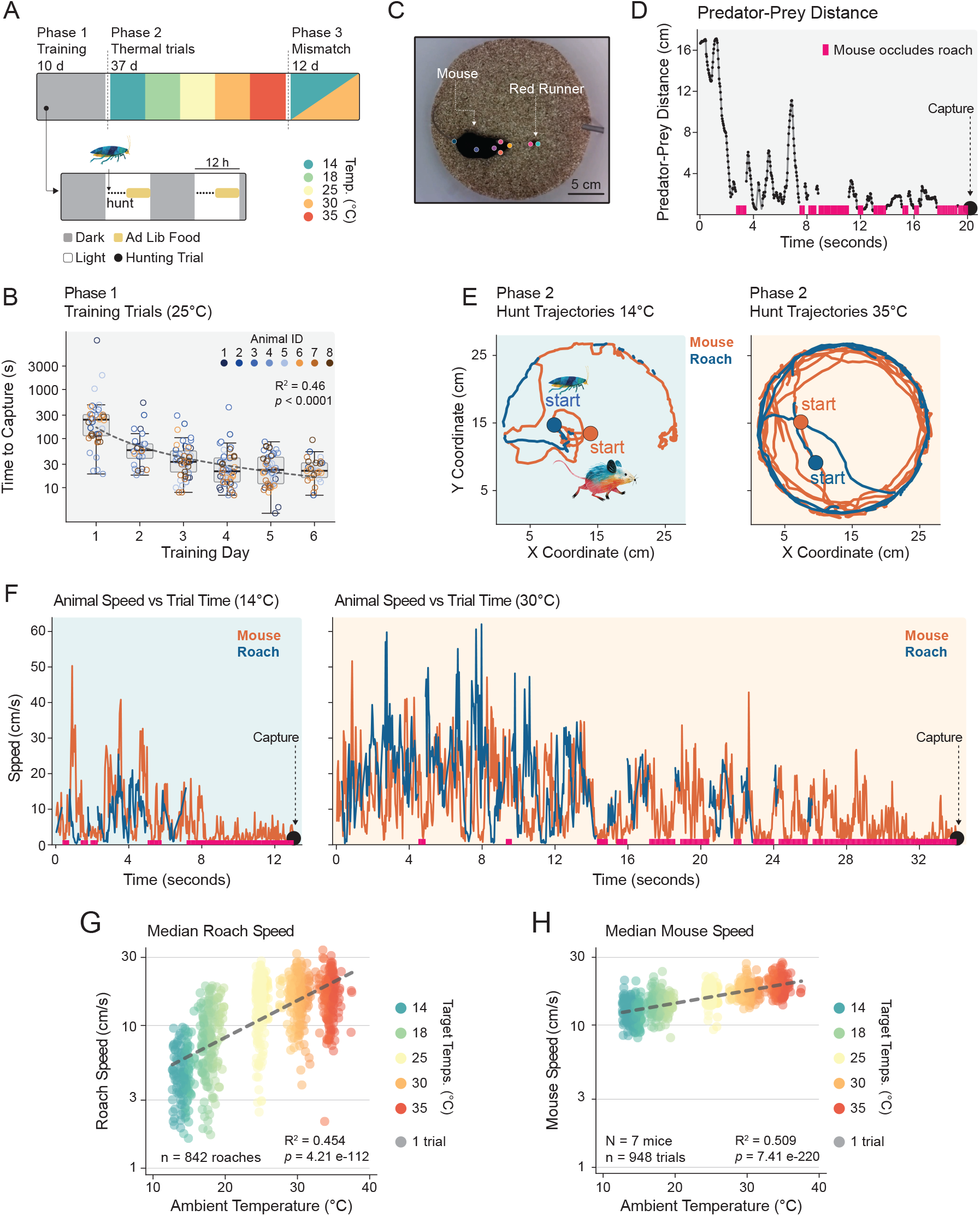
Mice rapidly learn to hunt and acutely match pursuit speed to temperature-dependent prey speed. **(A)** Schematic of overall experimental timeline (top) and daily trial protocol (bottom). *Top*: the experiment comprised three phases: Phase 1 ( Training, 10 d) at baseline room temperature (25*^◦^*C); Phase 2 (Thermal trials, 37 d), spanning five ambient temperatures (14*^◦^*C to 35 *^◦^*C; order counterbalanced across cohorts to control for temperature sequence effects); and Phase 3 (Mismatch trials, 12 d) featuring decoupled ambient and prey temperatures. *Bottom*: daily trial schedule showing light/dark cycles and food restriction timing. Each day consisted of overnight food restriction followed by six consecutive red runner cockroach hunting trials. **(B)** Time to capture across the first 6 consecutive days of initial room temperature (25*^◦^*C) training, shown grouped across animals (*n* = 8 mice, 30.3 *±* 2.3 trials per mouse, *n* = 242 total trials). Candidate learning models (linear, exponential, power-law) fit on log_10_-transformed capture times were evaluated using the Bayesian Information Criterion (BIC). Capture times decreased systematically across training, best described by a power-law fit (*y* = 170 *x^−^*^1.30^, *R*^2^ = 0.455, *p <* 0.001; ΔBIC *≥* 31.4) over exponential and linear models. Colored points denote individual trials per animal; box plots represent daily group medians and interquartile ranges (IQR). **(C)** Representative overhead frame from a single trial, shown with markerless pose estimates. **(D)** Predator-prey distance over the course of a representative trial, decreasing to capture. Pink bars indicate periods of contact between the mouse and red runner. **(E)** Mouse snout and red runner center trajectories from trial start to capture. *Left*, example cold (sluggish prey) trial at 14*^◦^*C. *Right*, example warm (fast, more active prey) trial at 30*^◦^*C. **(F)** Mouse spine and red runner center speed over trial time; capture marked at trial end. Same example cold (*left*) and warm (*right*) trials as in (**E**). Note the longer trial duration and greater peak velocities in the warm trial relative to the cold trial. **(G)** Red runner median speed during pursuit as a function of ambient temperature (*n* = 842 total trials). Each data point represents a single individual red runner. Log-transformed speeds were evaluated using a multiple OLS regression controlling for log-transformed body mass (*log*(*vel*) *∼ trial temp* +*log*(*mass*)). Red runner locomotor speed increased strongly with temperature (*R*^2^ = 0.454, *p <* 0.001), after controlling for body mass, which was also a statistically significant covariate (*p <* 0.001). **(H)** Mouse median trial speed as a function of ambient temperature (*n* = 7 mice, 135.4 *±* 13.2 trials per mouse, *n* = 948 total trials). Each data point represents a single individual trial. Log-transformed speeds were analyzed using an LMM with log-transformed red runner prey body mass included as a fixed covariate and animal ID as a random intercept (*log*(*vel*) *∼ trial temp* + *log*(*mass*) + (1*|animal*)). Mouse locomotor speed increased more modestly with ambient temperature (marginal *R_m_*^2^ = 0.509, *p <* 0.001), after controlling for prey body mass (*p <* 0.05), and accounting for baseline inter-individual variation across mice (random intercept, *p* = 0.05). Point color denotes mean trial temperature throughout; dashed lines are model fits rendered at median prey mass.

We reasoned that the simplest and most fundamental hunting strategy requires that mice acutely adjust their hunting behavior to the immediate speed of individual prey: faster red runners require faster pursuit. To test this, we next conducted hunting trials at different temperatures under the same protocol (**Fig. 2a**, top). During these trials, the effect of temperature on hunting dynamics was readily apparent: in cold trials, the sluggish prey were swiftly overtaken and subdued (**Fig. 2e,f**), whereas in warm trials, prey were markedly more active and challenging to capture (**Fig. 2e,f**). In line with these observations, red runner speed and acceleration during pursuit depended strongly on temperature (**Fig. 2g**; **Fig. S3a**), with movement rates increasing *∼* 4-fold until peaking at 30*^◦^*C (red runner pursuit speed: *R*^2^ = 0.454, *p <* 0.001; red runner pursuit acceleration: *R*^2^ = 0.307, *p <* 0.001). Consistent with the need to match the immediate demands of the trial, mouse speed and acceleration during pursuit also increased with temperature (**Fig. 2h**; **Fig. S3b**), but more modestly, as mice adjusted their movements to match that of the prey (mouse pursuit speed: *R*^2^ = 0.509, *p <* 0.001; mouse pursuit acceleration: *R*^2^ = 0.317, *p <* 0.001). Taken alongside the lack of temperature dependence in isolated mouse locomotion, these data demonstrate that mice flexibly adjust features of their hunting behaviors according to the difficulty of trials.

The simplest explanation for the adaptive matching of mouse speed to prey speed is direct adjustment of the innate, predatory behavioral program. This would comprise mice tuning pursuit kinetics in response to the immediate, observed prey. If so, the added difficulty in capturing faster prey should be accounted for entirely by pursuit dynamics. To assess this, we examined detailed features of hunting performance across temperatures. Consistent with the temperature-dependent changes in red runner locomotion, median capture time and total distance traveled increased *∼* 3-fold and 9-fold, respectively, from 14*^◦^*C to 30*^◦^*C (median capture time: *R*^2^ = 0.259, *p <* 0.001; median distance traveled: *R*^2^ = 0.437, *p <* 0.001) (**Fig. 3a,b**; **Fig. S4**; **Fig. S5**). Note that since red runner locomotion plateaus between 30 and 35*^◦^*C (**Fig. 2g; Fig. S3a**), features of hunting performance were only analyzed across the experimental temperature range wherein temperature predictably modifies red runner locomotion, i.e., from 14*^◦^*C to 30*^◦^*C. While at one level these results are unsurprising — faster prey should lead to longer and more extensive pursuits — any increase in capture time or distance traveled should, in principle, have been mitigated by the concomitant increase in mouse pursuit speeds observed at higher temperatures (**Fig. 3c**; median mouse speed: *R*^2^ = 0.465, *p <* 0.001). Because mice adjusted their pursuit speeds with temperature, predator-prey distance remained consistent across temperature conditions (**Fig. 3d**), with mice reliably maintaining a median pursuit distance of *∼* 4.2 cm (median predator-prey distance: *R*^2^ = 0.0124, *p* = 0.515). This temperature-invariant pursuit distance, in turn, mirrored a stable rate of contacts across temperatures (**Fig. 3e**), with mice tending to strike prey once every *∼* 3.7 seconds regardless of temperature (median strike rate: *R*^2^ = 0.00532, *p* = 0.105). Given that the pursuit distance and strike rate remained constant across temperatures, temperature-dependent increases in capture time and distance traveled must be driven by factors other than pursuit dynamics alone. One plausible explanation is that red runners were not only faster at warmer temperatures, but also more difficult to subdue upon contact. In other words, prey that are more metabolically active are better able to resist and escape capture, thus requiring a greater total number of contacts. Evidence supports this model: the number of strikes required to capture a red runner increased significantly with temperature, despite the fact that the rate of striking remained constant across conditions (**Fig. 3f**; median strikes per capture: *R*^2^ = 0.198, *p <* 0.001).

**Figure 3:**
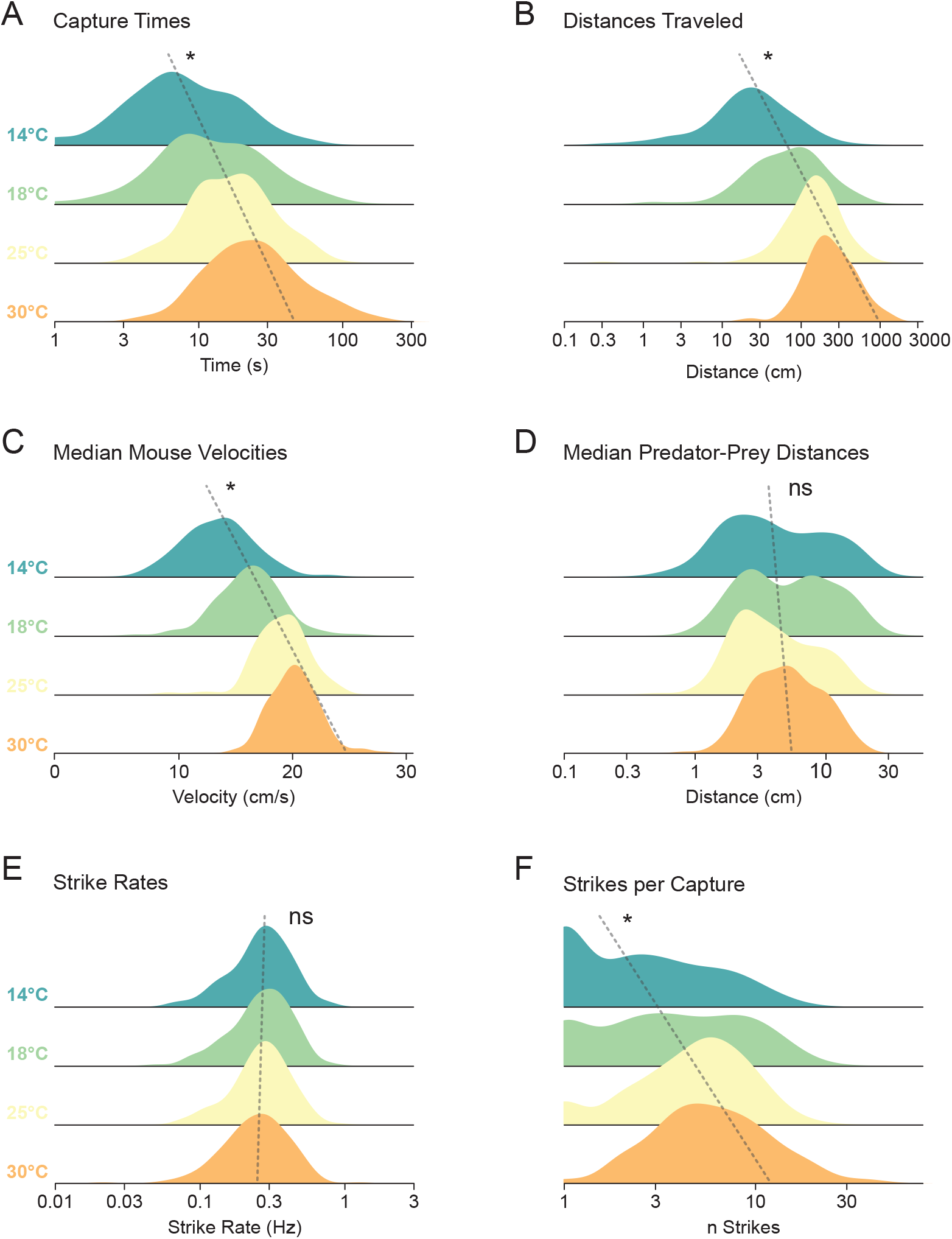
Behavior mitigation of effects of temperature on predation is feature-specific. Stacked kernel density estimation (KDE) curves (ridgeline plots) show the empirical probability distributions of hunting performance features across four ambient temperatures (14, 18, 25, 30*^◦^*C). Dashed lines indicate linear regressions fit through the coordinates of distribution medians. Directional shifts in central tendency across temperatures were evaluated using a two-tailed non-parametric permutation test (10,000 iterations) on log_10_-transformed group medians. **(A)** Capture time. Both central tendency and spread shifted toward higher values at warmer temperatures (*p <* 0.05). **(B)** Distance traveled per capture, which increased systematically with temperature (*p <* 0.05). **(C)** Median mouse speed, which increased systematically with temperature as mice matched faster prey (*p <* 0.05). **(D)** Median predator-prey distance, which remained consistent across temperatures (*p* = 0.294). **(E)** Strike rate, which remained consistent across temperatures (*p* = 0.233). **(F)** Strikes per capture, which increased systematically with temperature despite the constant strike rate (*p <* 0.05), consistent with warmer prey being more difficult to subdue upon contact.

Despite evidence for immediate tuning of predatory behavior, mice with experience hunting across a range of temperatures could, in theory, learn to treat temperature as a predictor of prey difficulty and use it to guide their hunting responses. This contrasts with the null hypothesis in which mice simply react to the moment-to-moment movements of their prey, despite having access to contextual environmental information with predictive value. To disambiguate these possibilities, we designed an experiment in which mice and red runners were maintained at *different* ambient temperatures prior to the hunting trial, thus violating potential expectations of symmetry between the ambient temperature of the arena and the speed of the red runner.

Until this point, mice and red runners were always acclimated to the same target temperature, across hundreds of trials spanning 14 to 35*^◦^*C (*∼* 40 per mouse per temp; **Fig. 2a**, top) — which provided ample opportunity for the mice to form reliable associations between thermal context and prey behavior. We then introduced a limited series of ‘temperature mismatch’ trials, in which the mouse and red runner were acclimated to *different* temperatures before the trial began. This allowed us to present warm prey (30*^◦^*C) to mice in cold arenas (14*^◦^*C), and conversely cold prey to mice in warm arenas, decoupling the mouse’s thermal context — and any associated expectations — from actual prey difficulty. If mice form and exploit such associations, violating them should disrupt hunting performance. If, instead, mouse predatory behavior is purely reactive, a temperature mismatch between predator and prey should have no effect; hunting success would depend solely on the prey’s actual speed, regardless of the mouse’s prior thermal context.

To avoid the possibility of disrupting any established associations between ambient temperature and red runner behavior, we conducted these mismatch trials infrequently — only once every other day, and only as the first of 6 daily trials, over a total of 12 days (**Fig. 4**). All other trials were run under standard, matched thermal conditions, both on mismatch days (trials 2-6) and alternating days with no mismatches (trials 1-6). As a result, in this phase of experiments, only 8.3% of trials comprised a mismatch, while the vast majority of trials (91.7%) reinforced any extant associations between red runner behavior and ambient temperature formed over the hundreds of prior trials. By this design, each mismatch was separated by 11 matched trials. Further, to control for the fact that mismatches were always the first trial, we compared mismatches only to the first trial of matched days, thus avoiding any possible confounds of short-term behavioral changes within a daily set of 6 trials. In total, we ran 72 trials per mouse (*n* = 7 mice) during the mismatch phase, of which 12 trials (6 mismatch, 6 matched; first of each day) were included for analysis (**Fig. 4**).

**Figure 4:**
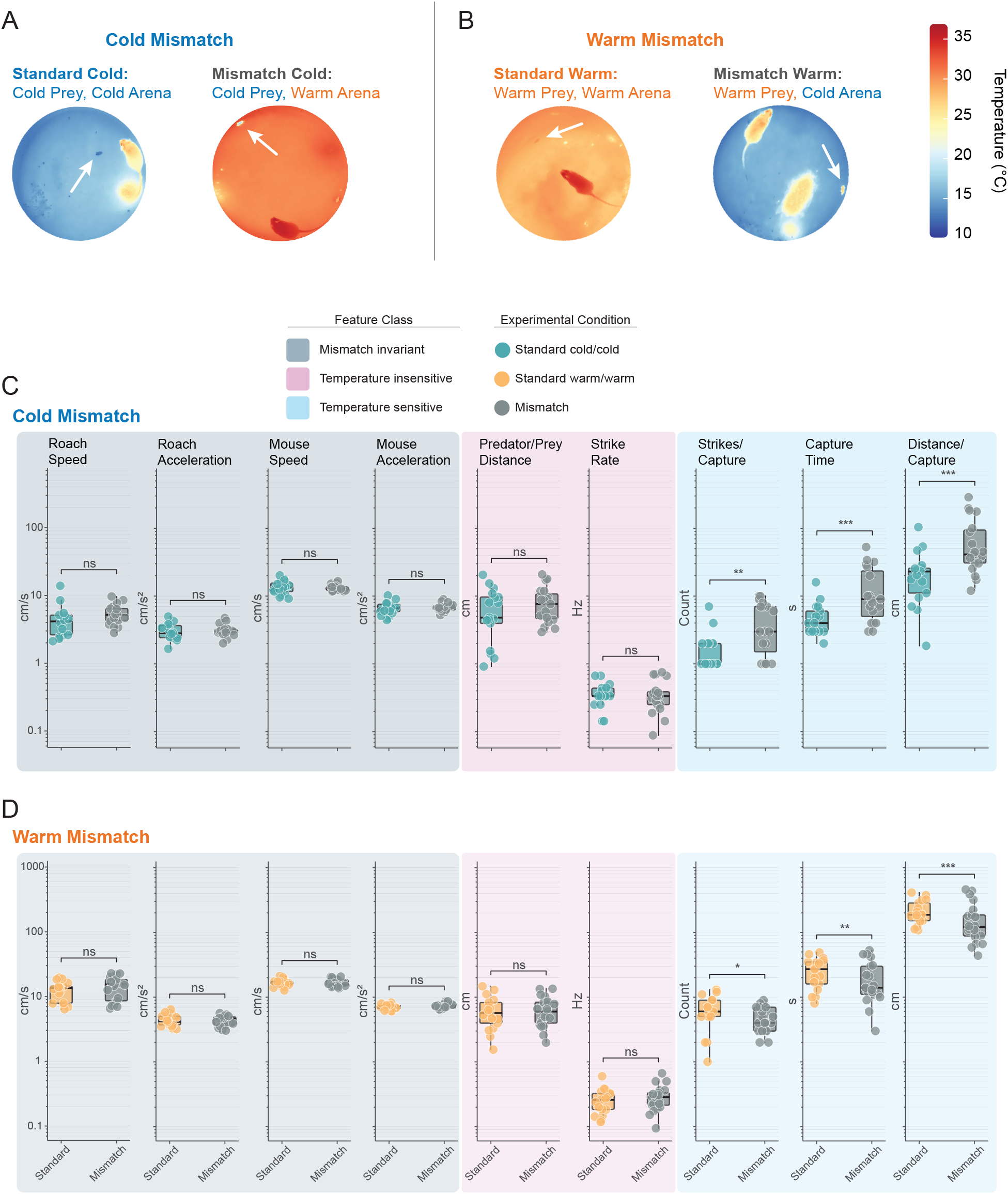
Temperature mismatch disrupts hunting performance, revealing learned use of predictive environmental context. Mismatch trials were run as the first of 6 daily trials, once every other day over 12 days, such that only 8.3% of trials were mismatches and each was separated by 11 matched trials; mismatches were compared only to the first trial of matched days (*n* = 7 mice). **(A)** Cold-mismatch design. Representative overhead thermal images of (Top) standard cold trial (14*^◦^*C) in which ambient temperature and red runner body temperature are both cold and (Bottom) mismatch cold trial in which the ambient temperature is warm (30*^◦^*C) while the red runner is cold. White arrow indicates location of the red runner. **(B)** Warm-mismatch design. Representative overhead thermal images of (Top) standard warm trial (30*^◦^*C) in which ambient temperature and red runner body temperature are both warm and (Bottom) mismatch warm trial in which the ambient temperature is cold (14*^◦^*C) while the red runner is warm. White arrow indicates location of the red runner. **(C)** Comparison of hunting performance features as a function of condition: standard cold vs. mismatch cold. Individual data points represent single trials and are shown on a logarithmic scale (box plots: median and IQR; points: individual trials). Movement features that do not differ between standard and mismatch trials are shaded in navy blue (i.e., ‘Mismatch invariant’). Behavioral features that were previously temperature insensitive are shaded in magenta, and those that were temperature sensitive are shaded in blue. **Prey performance**: red runner median speed and acceleration (*n* = 14 standard, *n* = 19 mismatch trials) were evaluated using multiple OLS regressions controlling for log-transformed body mass (*log*(*response*) *∼ trial cond* + *log*(*mass*)). Thermal mismatch did not significantly alter red runner speed (*p* = 0.093) or acceleration (*p* = 0.393). **Predator performance**: mouse median speed, acceleration, predator-prey distance, strike rate, strikes per capture, and distance traveled per capture (*n* = 17 standard, *n* = 19 mismatch trials) were all evaluated using LMMs with log-transformed red runner prey body mass included as a fixed covariate and animal ID as a random intercept (*log*(*response*) *∼ trial cond* + *log*(*mass*) +(1*|animal*)). Thermal mismatch did not significantly alter mouse motor performance (speed: *p* = 0.562; acceleration: *p* = 0.395) or spatial positioning features previously found to be temperature invariant (predator-prey distance: *p* = 0.509; strike rate: *p* = 0.973). In contrast, strikes per capture (*p <* 0.01, Cohen’s *d* = 1.22, marginal *R_m_*^2^ = 0.227), capture time (*p <* 0.001, *d* = 1.34, marginal *R_m_*^2^ = 0.257), and distance traveled per capture (*p <* 0.001, *d* = 1.33; marginal *R_m_*^2^ = 0.251) all increased significantly in mismatch trials. **(D)** Comparison of hunting performance features as a function of condition: standard warm (*n* = 17) vs. mismatch warm (*n* = 21 trials). Plotted and analyzed as in **(C)**. **Prey performance**: thermal mismatch did not significantly alter red runner speed (*p* = 0.766) or acceleration (*p* = 0.559). **Predator performance**: thermal mismatch did not significantly alter mouse motor performance (speed: *p* = 0.666; acceleration: *p* = 0.441) or spatial positioning features previously found to be temperature invariant (predator-prey distance: *p* = 0.753; strike rate: *p* = 0.188). In contrast, strikes per capture (*p <* 0.05, Cohen’s *d* = 0.70, marginal *R_m_*^2^ = 0.080), capture time (*p <* 0.01, *d* = 1.03, marginal *R_m_*^2^ = 0.144), and distance traveled per capture (*p <* 0.001, *d* = 1.44; marginal *R_m_*^2^ = 0.256) all decreased significantly in mismatch trials, opposite in direction to the cold-mismatch effects in **(C)**. Significance overlays: ns *p ≥* 0.05, * *p <* 0.05, ** *p <* 0.01, *** *p <* 0.001.

Given the relatively brief duration of a typical trial (ranging from 10.95 *±* 0.69 seconds at 14*^◦^*C to 32.59 *±* 2.16 seconds at 35*^◦^*C; values are mean±SEM), prey entering the arena at a mismatched temperature lacked sufficient time before being caught to thermally equilibrate to the ambient conditions, resulting in no meaningful alteration to body temperature during the trial (**Fig. S6**). Consistent with this, for a given red runner body temperature, average speed and acceleration did not differ significantly in matched and mismatched ambient conditions (**Fig. 4c,d**; both *p >* 0.05). Similarly, mouse speed and acceleration also did not change in mismatched trials compared to standard matched trials (**Fig. 4c,d**; both *p >* 0.05). Concretely, despite being in a warm arena, mice moved slowly when prey were cold and slow moving (and vice versa for the opposite temperature mismatch). These results demonstrate that mouse and red runner locomotion in mismatch and standard trials are unified by the temperature of the prey. Put another way, mice are able to match the speed of hot and fast-moving red runners equally well in both hot and cold environments. The null hypothesis, that mice acutely match their closed behavioral program to the locomotor rates of their prey, thus predicts that hunting outcomes should be equivalent between matched and mismatched temperature conditions. Alternatively, should predatory success differ in mismatched trials relative to matched, it would require a non-locomotor explanation.

Based on our data thus far (**Fig. 3**), if mice actively utilize ambient temperature to guide hunting behavior, such open-program learning should only be evident in hunting performance features that normally vary by temperature. In line with this logic, features that do not depend on ambient temperature, i.e., predator-prey distance and strike rate (**Fig. 3d,e**), should not differ between mismatched and matched trials. Consistent with this, we found that temperature mismatch did not alter predator-prey distance or strike rate (**Fig. 4c,d**; all *p >* 0.05). However, for all temperature-sensitive features (**Fig. 3a,b,f**), mismatch trials were significantly different than matched trials. For instance, for mice in warm arenas receiving cold prey (‘cold-mismatch’), strikes per capture, time to capture, and distance traveled per capture all increased significantly compared to standard conditions in which both the arena and prey were cold, with large effect sizes (**Fig. 4a,c**; all *p <* 0.01, Cohen’s d *>* 1.2). In ‘warm-mismatch’ conditions (cold arena, warm prey), time to capture and distance traveled per capture varied significantly compared to standard warm conditions, with medium to large effect sizes (**Fig. 4b,d**; both *p <* 0.01, Cohen’s d *>* 1.0), while strikes per capture was also significant (*p <* 0.05, Cohen’s d = 0.70). In the warm-mismatch experiment, the direction of the change was opposite of what we expected, with capture efficiency increasing in mismatch conditions relative to standard conditions.

Overall, mismatch trials significantly altered hunting outcomes even though statistics of mouse and red runner locomotion were unchanged, suggesting that environmental context prior to the trial influenced the subsequent expression of predatory hunting behavior. In cold-mismatch trials, capture performance was delayed relative to standard cold conditions (**Fig. 4a,c**), in accordance with our expectation that removing the predictive value of ambient temperature would impair hunting performance. Counterintuitively, in warm-mismatch trials, hunting efficiency increased (**Fig. 4b,d**). While the very fact of a consistent change supports the alternative hypothesis, the directionality of the change raises interesting questions about which costs mice aim to minimize via context-dependent hunting refinement.

## Discussion

In this study, we asked whether external environmental context shapes the expression of innate predatory hunting behavior. To test this, we conducted thousands of mouse/insect prey capture trials while systematically varying ambient temperature across a *∼* 20*^◦^*C range; because prey speed depends on temperature, this gave us direct control over the difficulty of the hunting task. Utilizing this novel approach, we trained naive laboratory mice to hunt red runner cockroaches at room temperature and then altered the ambient temperature between 14 and 35*^◦^*C to evaluate the resulting changes in hunting performance. Hunting performance depended strongly on ambient conditions, and outcomes were disrupted when the temperature experienced by the mouse differed from that of the prey. In these ‘temperature mismatch’ trials, in which the predictive value of the environmental context is removed, hunting performance was consistently altered. This suggests that mice learn to associate environmental cues with prey behavior and adjust their predatory behavior to anticipated demands. Our findings support a model in which innate, closed-program natural behaviors are refined through real-world experience and flexibly adjusted by top-down cognitive control. Our study establishes a powerful paradigm for linking ethologically relevant behaviors to informationally-rich environmental variables through the experimental control of ambient temperature.

Brains have evolved to effectively govern how animals participate in the world, ensuring individuals of a species are able to successfully navigate their unique ecological niche^11–14,62–66^. Natural behaviors arise when an animal — responding to the context of its environment — engages in stereotyped action patterns evolved to ensure its survival and reproduction^14–16^. Such behaviors, including courtship and mating, foraging and feeding, social communication and navigation, and predator avoidance and defense, are canonically viewed as products of innate, genetically-specified neural circuits that guarantee their reliable expression^67–73^. Yet although instinctive behaviors make up much of an animal’s natural repertoire, experience is often required to refine an inherent predisposition into a fully-developed, species-typical response. For example, most songbirds are born with an innate song template, and, even when raised without hearing other birds, will produce a rudimentary version of their species’ song. However, auditory exposure to a conspecific tutor and subsequent sensorimotor practice are needed to calibrate and refine this template into a fully developed adult song^74–76^. Similarly, even instinctive behaviors that appear fully functional the first time they are performed (i.e., ‘fixed action patterns’) can be modified by experience. In the classic case of the pecking response in gull chicks, newly hatched chicks will instinctively peck at a stimulus similar to a mother’s beak, such as a pointed object with a contrasting spot^77^. Yet subsequent work across multiple gull species has shown that visual discrimination of the parent improves over the first week of life: chicks initially peck at any elongated object but gradually come to preferentially peck at visual stimuli that more closely resemble the parent’s head^78^. This sharpening of perceptual selectivity, likely reinforced by successful feeding, together with improvements in pecking accuracy, suggests that even natural behaviors traditionally regarded as solely instinctive can undergo experience-dependent refinement. However, each of these examples — and the rapid performance gains mice show in our own initial hunting trials (**Fig. 2b**) — involves the refinement of an instinctive behavior through immediate, direct feedback on the behavior itself. A chick pecks and is fed; a mouse strikes and captures prey. In each case the reinforcing signal bears directly on the action being refined. None of these cases addresses whether an instinctive behavior can be modified through *complex learning*, in which an indirect environmental cue that carries no direct sensory information about the action is used to anticipate task demands and select among strategies in advance. This form of adaptation, rather than mere refinement, has typically been regarded as a hallmark of flexible, top-down cognition^79–82^. In the context of predatory hunting behavior, performance is instead generally attributed to innate circuitry tuned to each species’ unique ecological needs^6,8,43,45^. Yet despite its clear ecological importance, the extent to which natural hunting behavior is adjusted according to real-world experience has remained largely unexplored^51^.

In this work, we introduced a thermal mismatch to test whether mice learn to associate indirect environmental variables with the task at hand and proactively adjust their behavioral strategies. Our results show that disrupting the ambient temperature cues that typically signal prey difficulty reliably altered hunting performance. This finding suggests that, with experience, mice integrate contextually relevant information into their predatory behavior and use it to forecast the demands of a hunt and modulate their actions accordingly. The capacity to adjust instinctive behaviors to local conditions is advantageous because natural habitats are complex, variable, and often unpredictable^83,84^. Behavioral flexibility therefore enables animals to respond more effectively to shifting ecological pressures, supporting short-term performance while potentially facilitating longer-term niche expansion over evolutionary timescales^65,85,86^. While breaking an established contingency can impair task performance, the magnitude of that cost depends on how strongly the contextual cue influences the underlying behavior. In our experiments, temperature mismatch consistently altered hunting efficiency (as measured by time to capture, distance to capture, and strikes per capture), yet mice continued to deploy the stereotyped search-pursue-bite sequence successfully. Temperature cues thus refine execution but do not reorganize the innate hunting program: although mice had formed associations between ambient temperature and prey difficulty, they still hunted proficiently when those expectations were violated because the underlying hunting strategy was largely invariant across temperatures. This may reflect the simplicity of our assay — a bare, circular arena affords mice few opportunities to modify or elaborate their instinctive hunting tactics. In a more heterogeneous environment containing obstacles, variable terrain, and potential prey refuges, mice would have greater latitude to apply learning and cognition to the hunting challenges such a habitat imposes^87^. Under such conditions, in which the hunting strategy itself must adapt to complex environmental structure^88^, temperature cues might carry even greater predictive value for anticipating prey difficulty and selecting among behavioral options. Even in our structurally simple arena, however, thermal mismatch altered every temperature-sensitive performance metric we quantified. Mice therefore form contextual strategies based on ambient temperature even under conditions that limit their options for acting on them.

An alternative explanation for the mismatch results is that arena temperature acted directly on the physiological state of the mouse, for example by increasing energetic demand and appetitive drive in cold arenas, producing performance differences without any learned association. Two features of the design argue against this account. Although acute cold exposure can increase feeding drive within minutes in ad libitum-fed mice^89^, every trial in our study followed 14-16 h of food restriction — a fasted state that strongly activates the same hypothalamic hunger circuitry^90^ and leaves little room for a 30-minute thermal exposure to further raise appetitive drive. A motivational effect should also appear in pursuit vigor, yet mouse speed, acceleration, and strike rate were unchanged in mismatch trials (**Fig. 4c,d**); only outcome measures shifted. These kinematic measures do not fully capture persistence during prey subdual, but a motivational account that alters subdual while leaving pursuit untouched would require an additional assumption for which we see no evidence.

The apparent paradox of increased hunting efficiency during the warm-mismatch condition (**Fig. 4b,d**) suggests a misalignment between our experimental metrics and the animal’s internal optimization criteria. While we define efficiency using external parameters like capture time, total distance, and strike count, the mouse may be optimizing for a completely different objective function, such as minimizing cumulative energy expenditure. Thus, though our data demonstrate that ambient temperature shapes the mouse’s predatory behavior, the nature of this optimization remains to be characterized. Speculatively, under standard warm conditions, where prey are relatively fast and difficult to capture, mice may adopt a conservative hunting strategy that balances physical effort against expected payoff. In a simple, closed arena with no escape routes, small reductions in capture time may not justify the added energetic cost. Under this framework, when a mouse expects slow, cold prey, it may instead opt for a high-vigor, sprint-like pursuit, because the anticipated brevity of the hunt keeps total energetic cost low. In a warm-mismatch trial, this expectation of an easy, rapid catch leads the mouse to pursue fast, warm prey with greater intensity, which improves its measured efficiency. The same model explains the cold-mismatch results: a mouse that expects fast, warm prey and paces its effort to conserve energy will perform more slowly against slow, cold prey. Directly testing this interpretation would require real-time metabolic measurements, but it is consistent with optimal foraging theory^91–95^, in which predators calibrate effort to anticipated prey difficulty.

Beyond the scale of the individual organism, these findings connect to temperature’s role in structuring global ecosystems. An individual predator’s capacity to exploit temperature as a predictive cue is limited by the degrees of freedom in its immediate environment, but at macroecological scales, temperature structures entire food webs through its influence on ectotherm metabolism^96,97^. Ectothermic and endothermic animals are tightly integrated across global food webs, with ambient temperature mediating their trophic interactions^98,99^. Because cold temperatures depress ectotherm locomotor performance while leaving endotherms relatively unaffected, endothermic predators are favored where prey are cold and sluggish, whereas ectothermic predators are favored in warmer environments^100,101^. By isolating a single predator-prey pair and experimentally dissociating thermal context from prey speed, our study adds a cognitive and behavioral dimension to these macroecological patterns^88^: temperature constrains hunting outcomes, and an endothermic predator can learn to anticipate those constraints. The global thermal structuring of predator-prey interactions may therefore reflect not only the passive biophysics of prey metabolism but also the capacity of predators to learn and exploit the thermal regularities of their niche.

## Acknowledgments

This project was supported by the National Institutes of Health (NIH) BRAIN Initiative award R01NS118442 (K.B.H.), National Science Foundation (NSF) Research Traineeship award 2152221 (J.L.A.), and NIH Institutional National Research Service awards T32EY013360 from the National Eye Institute and T32NS121881 from the National Institute of Neurological Disorders and Stroke (J.L.A.). Additional support provided by the Living Earth Collaborative at Washington University (J.M.G.) and NSF award DEB-2017740 (A.I.D.). J.M.G. and A.I.D. were also supported by NSF award DEB-1838346. We thank James N. McGregor for helpful suggestions, and Bryan Higashikubo, Marcos Bermudez and Luke Long for assistance with experimental procedures.

## Author Contributions

J.L.A., J.M.G., A.I.D., and K.B.H. designed the experiments. J.L.A. and J.M.G. performed experiments. J.L.A., J.M.G., K.B.N., V.S., and S.J.B. analyzed data. J.L.A. and K.B.H. wrote the manuscript. All authors reviewed the manuscript and contributed to improvements.

## Data Availability

Experimental data is available upon request.

## Code Availability

All code for analyses is available at https://github.com/hengenlab

## Methods

### Experimental model and subject details

All procedures were conducted in accordance with protocols approved by the Washington University in Saint Louis Institutional Animal Care and Use Committee (IACUC), following NIH guidelines for the care and use of research animals. Eight C57BL/6J mice (five female, three male) were used in this study (Supplier: The Jackson Laboratory). One male mouse ceased hunting after the initial room temperature trials and was excluded from the subsequent thermal trials (see below). Mice were seventeen weeks old at the time of prey capture experiments. No differences in predation were observed between males and females and so their data were pooled. Turkestan red runner cockroaches (Supplier: Caribbean Mealworms) aged three to four months and weighing 0.05 - 0.20 g (mean = 0.1 g), were used as prey for all capture experiments.

### Prey capture experiments

Mice were singly housed in circular hunting arenas (36.83 cm H *×* 25.87 cm D) beginning forty-eight hours before hunting trials commenced, with food pellets and water available *ad libitum*. These arenas doubled as home cages for the duration of the experiments and sat within ventilated, temperature-controlled chambers on a 12:12 light/dark cycle. Once the mice had habituated to the enclosures for forty-eight hours, three red runners were placed in each room-temperature arena (25*^◦^*C) overnight (i.e., the 12 h dark period), alongside *ad libitum* access to food pellets. Red runners still present at the close of the 12 h dark period were removed. This procedure was repeated the following day, yielding 2 d of nocturnal red runner exposure in total.

### Initial Trials

Ten days of initial hunting trials at room temperature followed (**Fig. 2a,b**). A trial began with the introduction of a single red runner to the arena, and mouse-red runner interactions were captured on an overhead camera (30 fps; e3Vision; White Matter). On each day of trials, up to six red runners were sequentially introduced to each mouse (*≈* 60 trials per mouse in total), with 10 minutes separating consecutive trials. A trial ended once the red runner was incapacitated. Food pellets were removed 14-16 h before each day of trials, which began 1 h after light onset, and were returned to the arenas for 8-10 h at the end of each trial day.

### Thermal Trials

Thermal trials at 14, 18, 25 (room temperature), 30, and 35*^◦^*C were conducted after the 10 d of initial trials (these trials are also analyzed in Grady et al., 2026^102^, which tests thermal-asymmetry predictions in an ecological context). Protocols were the same, with the addition of ambient temperature control. Mice and red runners shared a single temperature-controlled chamber, so that both experienced identical thermal conditions, and chambers were held at the target temperature for 30 minutes before trials began to allow red runner body temperature to equilibrate passively to ambient conditions (**Fig. 1d**). Thermal trials proceeded in two phases. The first ran for 27 d, with the target temperature changed every 5 d (*≈* 30 trials per temperature per mouse). Order effects were controlled by exposing half the mice to cold before warm conditions and half to warm before cold. To verify that temperature-dependent performance persisted after weeks of training, a second phase of thermal trials was run for 10 d, with the target temperature changed daily (*≈* 12 trials per temperature per mouse). No differences in predation emerged between phases, and all data were therefore pooled for analysis. Trials were discarded when mice displayed signs of overheating at 35*^◦^*C (e.g., splayed legs), hunted without persistence, declined to eat prey, or when recording errors occurred (13.1% of trials in total).

### Thermal Mismatch Trials

Thermal mismatch trials were conducted following the 37 d of standard thermal trials and were designed to test whether mice exploit temperature cues to anticipate prey behavior. In contrast to standard thermal trials in which the red runners and mice were maintained at matching target temperatures, thermal mismatch trials involved maintaining red runners at a different temperature than the mouse arenas to which they were subsequently introduced. Specifically, cold red runners (cooled to 14 *^◦^*C) were placed into warm mouse arenas (heated to 30 *^◦^*C) and vice versa (**Fig. 4a,b**). The thermal mismatch experiments were conducted for 12 d. Half the mice began in cold conditions (14*^◦^*C) and half in warm (30*^◦^*C), with conditions reversed after 6 days. Across each set of 6 d, standard (i.e., matched) thermal trials were carried out as previously described (i.e., 6 trials per mouse per day), except every other day in which a thermal mismatch trial was substituted for trial no. 1 and followed by five trials in matched conditions (yielding 3 mismatch trials total per mouse across each set of 6 d). Overall, each of the seven mice underwent 6 mismatch trials, for 42 mismatch trials in total, with each mismatch trial separated from the next by 11 trials.

For analysis, locomotion and hunting performance for mismatch trials were compared to that of the first trial of matched days, to control for any daily order effect. Specifically, mice hunting warm red runners in warm arenas (‘standard warm’) were compared with mice hunting warm red runners in cold arenas (‘mismatch warm’), and the equivalent comparison was made between ‘standard cold’ and ‘mismatch cold’ conditions. Thermally mismatched red runners did eventually equilibrate toward ambient temperature, but they were typically consumed within 3 to 20 s, which limited the drift in red runner body temperature (**Fig. S6**).

### Movement tracking and analysis

Videos of mice and red runners were recorded with an overhead camera (30 fps; e3Vision; White Matter). Time to capture was defined as the time between trial start (i.e., when the red runner enters the arena) and capture (i.e., when the red runner is decapitated). Time to capture was scored manually for every trial, and scoring accuracy was checked on a subset of videos by two trained scorers. Markerless pose estimation on both mouse and red runner was performed with DeepLabCut^61^ (**Fig. 2c**), tracking six anatomical points on the mouse (snout, left ear, right ear, shoulder, spine, tail base) and two on the red runner (front and back). Additional points, such as the red runner center, were derived from these six by midpoint calculation,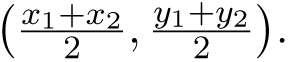 Second-order features were then computed for each animal with a custom Python script: distance traveled 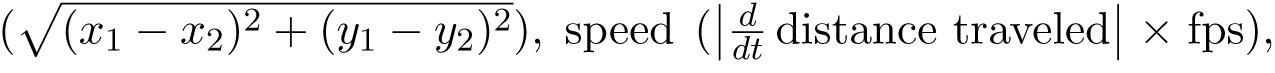 and acceleration 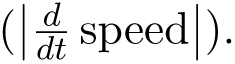 A strike was scored whenever the mouse occluded both red runner points for 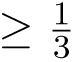 (**Fig. 2d**). Red runner movement was calculated with respect to the red runner center and mouse movement was calculated with respect to the mouse spine.

### Video post-processing

Videos were recorded at 30 fps and markerless pose estimation was conducted on the mouse and red runner using DeepLabCut^61^ across 9 million total frames. Because the model occasionally returned erroneous pose estimates, a series of post-processing steps was conducted to identify and remove outliers (custom Python script). For each video, a frequency distribution of frame-to-frame changes in spatial position was inspected for outliers and compared these against the corresponding video segment in order to set a conservative exclusion threshold. Pose estimates were discarded whenever the distance traveled between consecutive frames exceeded 3 cm (equivalent to 90 cm/s). Additionally, red runner pose estimates were discarded whenever the front and back values differed by more than 2 cm (i.e., more than one body length).

Next, movement values were excluded that did not reflect genuine locomotion, such as grooming or pixel jitter. This was defined as the 95% quantile of speed and acceleration for mice or red runners that were visually confirmed to be stationary in videos (**Fig. S7**). These values were taken as the threshold separating non-locomotory from locomotory movement; values falling below it were excluded from analysis. Any remaining outliers were identified and removed using the following outlier formula: *Q*2 *± C × IQR*, where *Q*2 is the median, *IQR* the 25–75% inter-quartile range, and *C* a coefficient (note that *C* = 1.5 defines the outlier range of a conventional box plot, corresponding to 2.7 standard deviations in a normally distributed dataset). For a given trial, *C* = 3 (equivalent to 4.5 standard deviations) was used, and for pooled red runners or mice at each thermal condition, *C* = 6 (equivalent to 9 standard deviations) was used.

### Temperature recordings

To measure brain temperature, mice were placed in temperature-controlled chambers (14*^◦^*C, *n* = 4; 30*^◦^*C, *n* = 4) for 30 minutes and then rapidly anesthetized with 5% isoflurane. Once loss of consciousness was confirmed (*≈* 10 s), each mouse was decapitated and a sterile temperature probe was inserted immediately into the brainstem via the foramen magnum. Temperature was sampled for 5 s at 1 Hz with a Leaton Digital Dual Thermometer (**Fig. 1c**).

To measure insect internal temperature, red runners were placed in temperature-controlled chambers (14*^◦^*C, *n* = 4; 18*^◦^*C, *n* = 2; 25 °C, *n* = 4; 30 °C, *n* = 5) for 30 minutes, after which a sterile temperature probe was inserted into the abdomen of each insect. Temperature was sampled for 5 s at 1 Hz with a Leaton Digital Dual Thermometer (**Fig. 1d**).

For thermal imaging, naive mice and red runners were placed in temperature-controlled chambers cooled to 14*^◦^*C or heated to 30*^◦^*C. After 30 minutes, one mouse and one red runner drawn from matched thermal conditions (i.e., 14*^◦^*C for both or 30*^◦^*C for both) were paired and imaged with a thermal camera (FLIR E53; Teledyne FLIR). The same imaging protocol was repeated with mice and red runners drawn from mismatched thermal conditions (**Fig. 4a,b**).

## Statistical analysis

All analyses were performed in Python using custom scripts. Full details are available in the code (see **Code Availability**).

Body temperature as a function of ambient temperature (**Fig. 1c,d**) was modeled using ordinary least squares (OLS) linear regression (*body temp ∼ trial temp*), with slope significance evaluated using a two-tailed Student’s t-test (*df* = *n −* 2). All measures of red runner movement — both solitary (**Fig. 1f**; **Fig. S1b**) and during pursuit (**Fig. 2g**; **Fig. S3a**; **Fig. 4c,d**) — were modeled using multiple OLS linear regression (*log*(*response*) *∼ trial temp* + *log*(*mass*)). Because each red runner was tested in a single trial, individual observations were statistically independent. In these models, both the dependent variable (median movement rate) and red runner body mass were natural log-transformed (ln) to satisfy model assumptions of normality and homoscedasticity, while controlling for size-dependent allometric scaling effects (**Fig. S8**). The statistical significance of the temperature effect was determined using a two-tailed Student’s t-test on the regression coefficient (*df* = *n −* 3, adjusting for the prey mass covariate). Reported *R*^2^ values represent the OLS coefficient of determination, and plotted trendlines reflect back-transformed predictions with mass held constant at the sample median.

All analyses of mouse movement (**Fig. 1e**; **Fig. 2h**; **Fig. S1a**; **Fig. S3b**), hunting-outcome features (**Fig. 3**), and mismatch comparisons (**Fig. 4c,d**) were performed using linear mixed-effects models (LMMs) with a random intercept per animal, to account for non-independence arising from repeated measures across individual mice. Models included continuous ambient temperature (*trial temp*) and prey body mass (*mass*) as fixed predictors, alongside a random intercept per animal (*log*(*response*) *∼ trial temp* + *log*(*mass*) + (1*|animal*)). Response variables and prey mass were natural log-transformed (ln) prior to fitting to satisfy model assumptions of normality and homoscedasticity, while controlling for size-dependent allometric scaling effects. Statistical significance of fixed-effect terms was evaluated using two-tailed Wald z-tests. The statistical contribution of the random intercept (animal ID) was evaluated via a Likelihood Ratio Test (LRT) comparing the LMM against an unclustered single-level multiple OLS regression, applying a 50:50 mixture chi-squared boundary correction 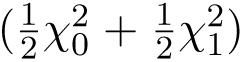 to account for testing a variance component at its boundary (σ_α_^2^ = 0). Goodness-of-fit was quantified following Nakagawa and Schielzeth, reporting both Marginal *R*^2^ (*R_m_*^2^; variance explained by fixed effects alone) and Conditional *R*^2^ (*R_c_*^2^; variance explained by fixed and random effects combined). Plotted trendlines represent back-transformed population-level fixed-effects predictions across trial temperatures, evaluated with prey mass held constant at the sample median.

To assess the statistical significance of the directional shifts in hunting performance features across temperature groups (**Fig. 3**), a two-tailed non-parametric permutation test (*n* = 10, 000 permutations) was performed. For each permutation, the temperature group labels defining the stacked kernel density estimation (KDE) distributions were shuffled across trials, and null group medians were recalculated in log_10_ space. An OLS linear regression was fit through these null medians as a function of temperature to construct an empirical null distribution of slopes. The true observed regression slope was then evaluated against this null distribution to calculate a two-tailed p-value. This statistical approach directly tests the directional shift in central tendency tracked visually by the dashed trendlines in the ridgeline plots.

For the mismatch comparisons (**Fig. 4c,d**), which were modeled using multiple OLS and LMMs as detailed above, effect sizes were computed as Cohen’s *d*. Here, *d* was defined as the model-adjusted difference in estimated marginal means divided by the pooled standard deviation of the model residuals. Confidence intervals for *d* were obtained via normal approximation, with the standard error derived from group sample sizes and the effect magnitude.

## Supplemental Figures

**Figure S1:**
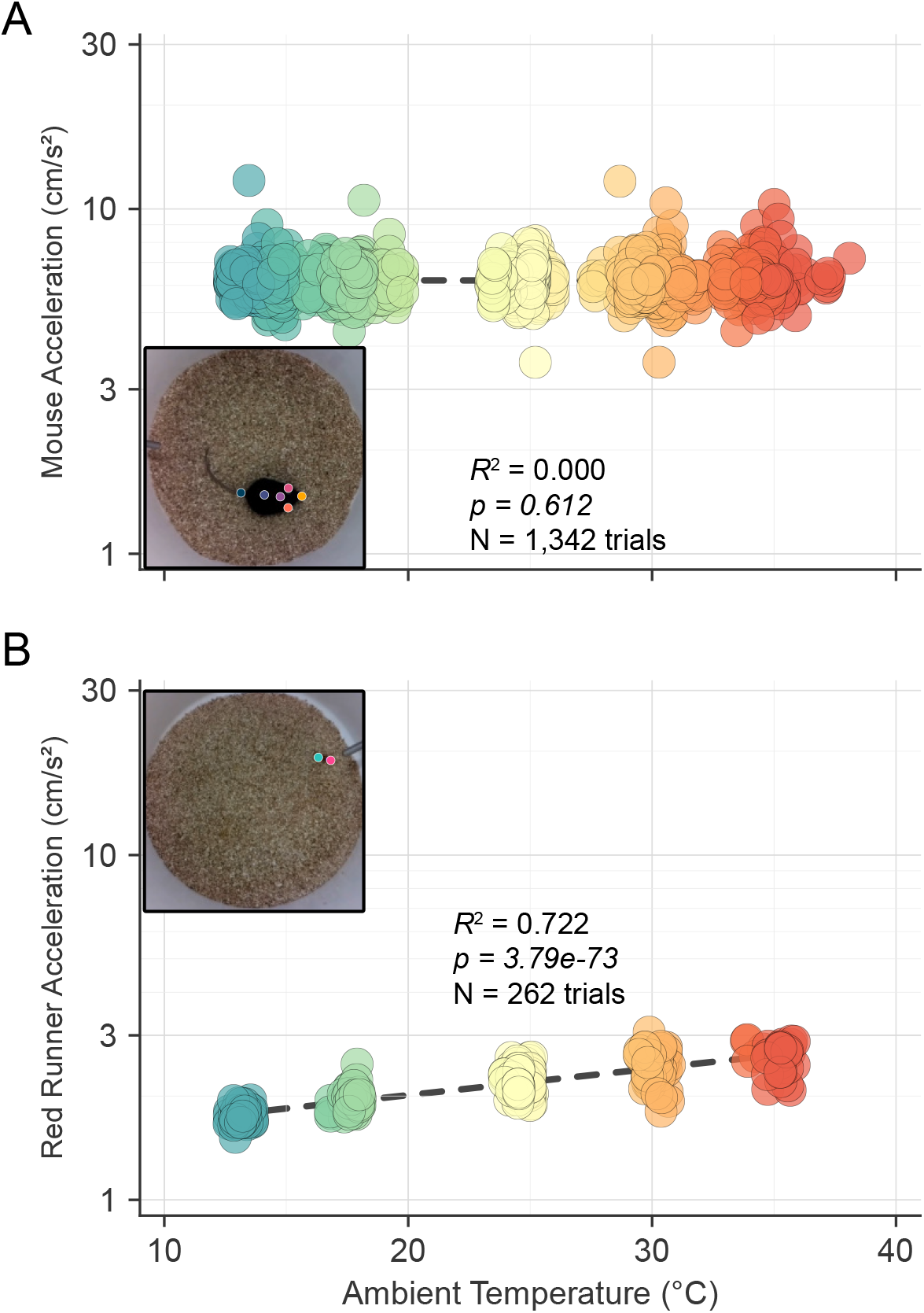
Locomotor acceleration in isolation is temperature-invariant in mice but temperature-dependent in red runners. Median locomotor acceleration as a function of ambient temperature for solitary animals. Each marker is a single trial, colored by the actual mean trial temperature; dashed lines are model fits. **(A)** Mouse acceleration assessed in isolation (*n* = 7 mice, 191.7 *±* 15.7 trials per mouse, *n* = 1,342 total trials). Log-transformed accelerations were analyzed using a Linear Mixed-Effects Model (LMM) with animal ID as a random intercept (*log*(*acc*) *∼ trial temp* + (1*|animal*)). Mouse acceleration was invariant to ambient temperature (marginal *R_m_*^2^ = 0.000, *p* = 0.612), after accounting for significant baseline differences between individual animals (random intercept *p <* 0.001). **(B)** Red runner acceleration assessed in isolation (*n* = 262 trials). Each data point represents a single individual red runner. Log-transformed accelerations were evaluated using a multiple OLS regression controlling for log-transformed body mass (*log*(*acc*) *∼ trial temp* + *log*(*mass*)). Red runner locomotor acceleration increased significantly with temperature (*R*^2^ = 0.722, *p <* 0.001), after controlling for body mass, which was not a statistically significant predictor (*p* = 0.088; fit line rendered at median prey mass). Insets show representative overhead video frames with markerless pose estimates.

**Figure S2:**
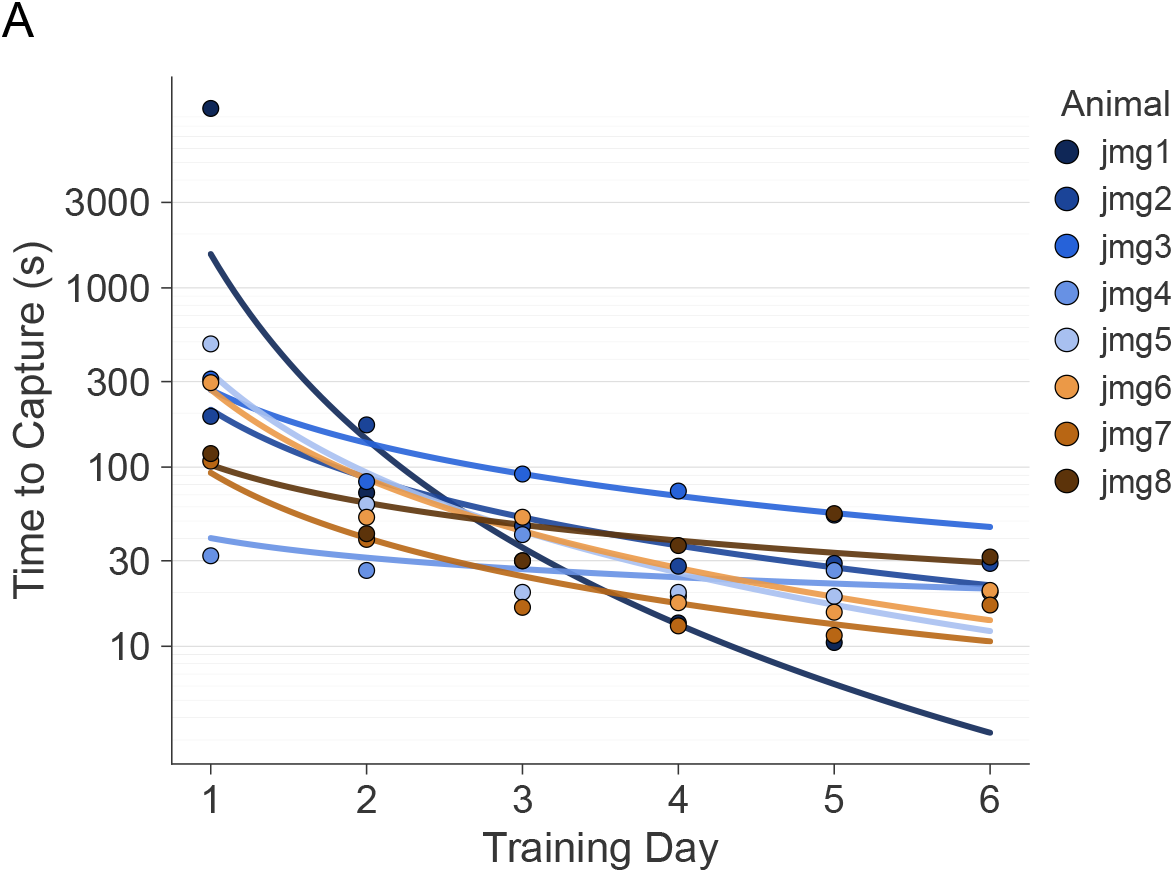
Per-animal room temperature learning curves. **(A)** Time to capture as a function of training day for each individual mouse (jmg1–jmg8; *n* = 8) during the initial room temperature (25*^◦^*C) training period. Points are per-day median capture times for each animal; lines are per-animal power-law fits selected by BIC (log_10_-time space). Note that for one animal (jmg1), an extremely high initial capture time on Day 1 led to a steeper power-law fit relative to other animals.

**Figure S3:**
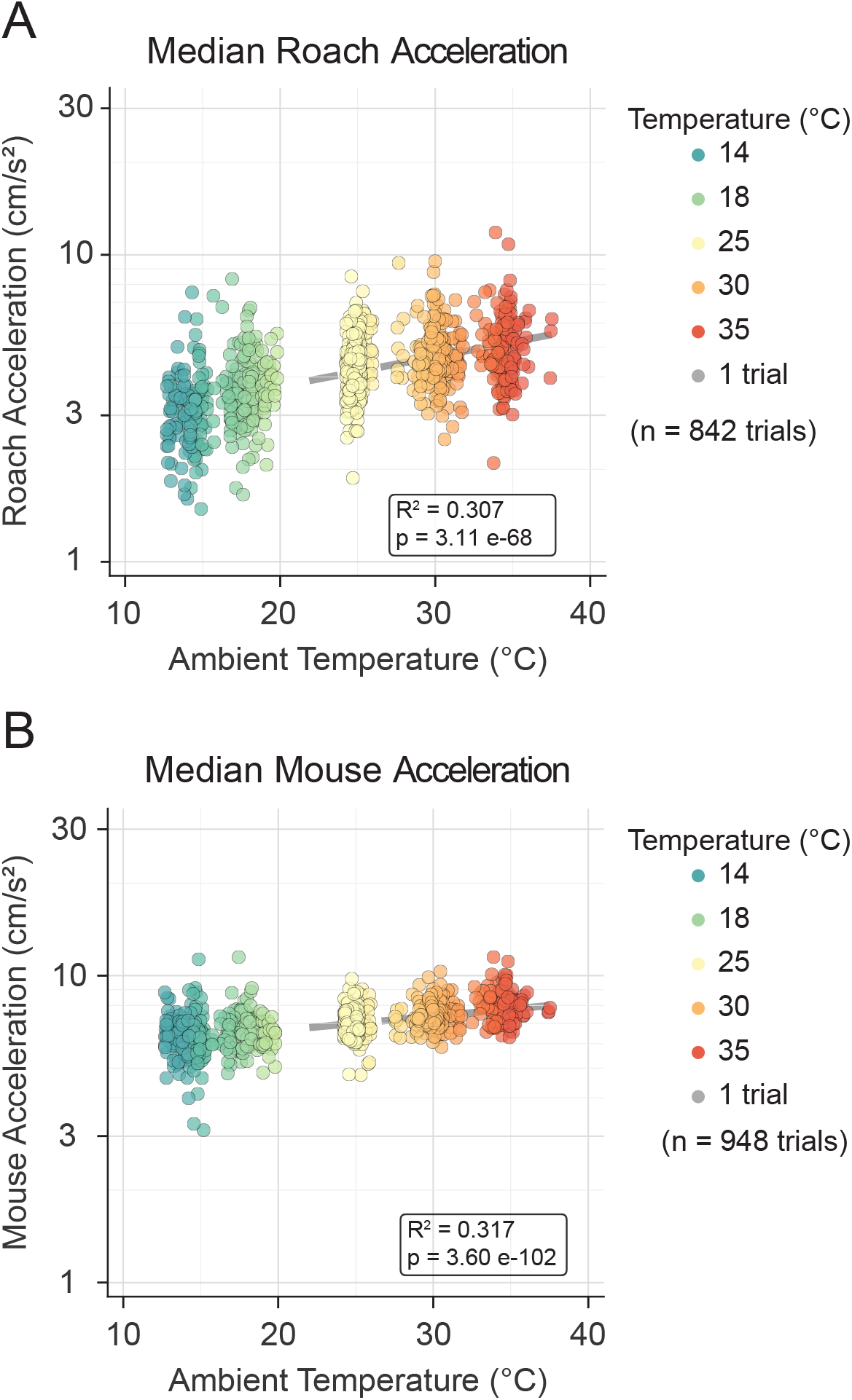
During pursuit, both red runner and mouse acceleration increase with ambient temperature. Median locomotor acceleration during hunting trials as a function of ambient temperature. Each marker is a single trial, colored by the actual mean trial temperature; dashed lines are model fits rendered at median prey mass. **(A)** Red runner acceleration assessed in isolation (*n* = 842 trials). Each data point represents a single individual red runner. Log-transformed accelerations were evaluated using a multiple OLS regression controlling for log-transformed body mass (*log*(*acc*) *∼ trial temp* + *log*(*mass*)). Red runner locomotor acceleration increased significantly with temperature (*R*^2^ = 0.308, *p <* 0.001), after controlling for body mass, which was also a statistically significant covariate (*p <* 0.001). **(B)** Mouse acceleration assessed in isolation (*n* = 7 mice, 135.4 *±* 13.2 trials per mouse, *n* = 948 total trials). Each data point represents a single individual trial. Log-transformed accelerations were analyzed using an LMM with log-transformed red runner prey body mass included as a fixed covariate and animal ID as a random intercept (*log*(*acc*) *∼ trial temp* + *log*(*mass*) + (1*|animal*)). Mouse locomotor acceleration increased more modestly with ambient temperature (marginal *R_m_*^2^ = 0.317, *p <* 0.001), after controlling for prey body mass (*p <* 0.05), and accounting for baseline inter-individual variation across mice (random intercept *p <* 0.05).

**Figure S4:**
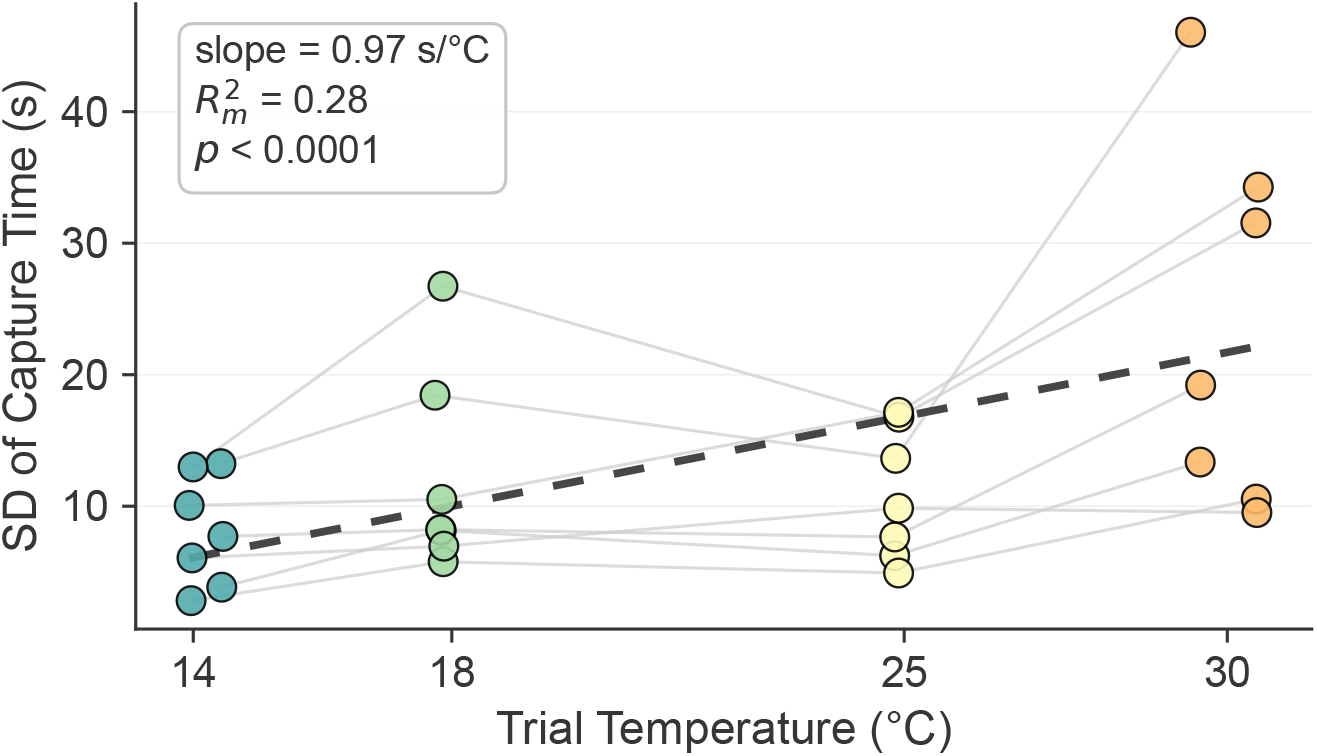
The spread of capture times increases with ambient temperature. Standard deviation (SD) of capture time (seconds) plotted against mean trial temperature for each mouse across target temperature conditions (*n* = 7 mice, 28 per-mouse summary observations). Points represent individual per-mouse SD values within each target temperature group, colored by target temperature; faint gray lines connect observations from the same individual across temperatures. The dashed black line represents the population-level trendline from an additive linear mixed-effects model (LMM) evaluated at the overall mean log-transformed prey mass (*SD ∼ mean trial temp* + *mean log mass* + 1*|animal*). Capture time variability significantly increased with trial temperature (*p <* 0.001); slope = 0.97s/*^◦^*C), after accounting for mean log prey mass (*p <* 0.05) and significant baseline inter-individual variability across mice (random intercept, *p <* 0.001; marginal *R_m_*^2^ = 0.278, conditional *R_c_*^2^ = 0.670).

**Figure S5:**
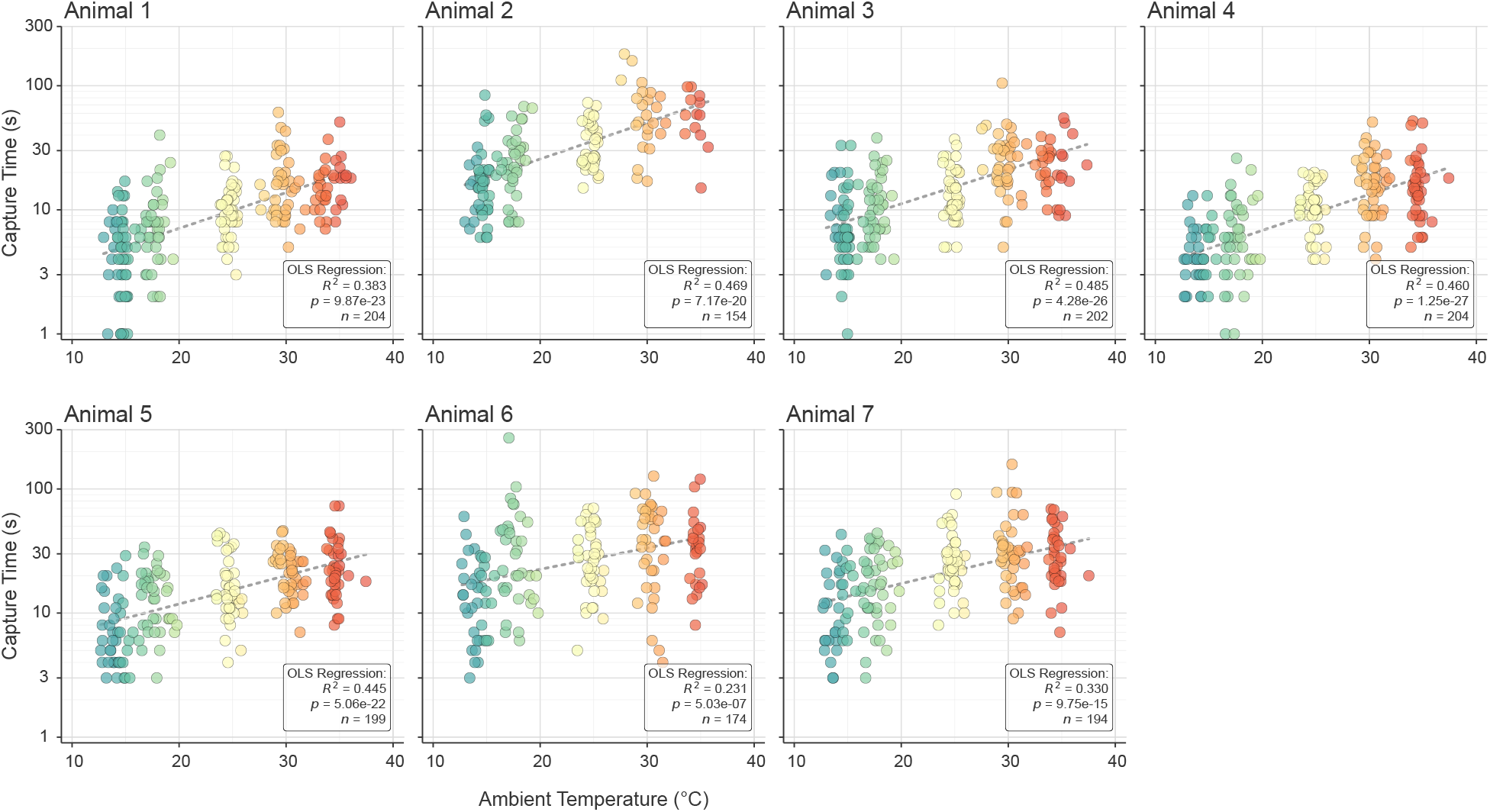
Individual mice thermal hunting trials. Time to capture during thermal hunting trials as a function of ambient temperature. Individual panels display performance metrics grouped by animal. Individual markers represent single trials colored by target temperature. Dashed lines are model fits shown at median red runner mass. Log-transformed capture times were evaluated using multiple OLS regression controlling for log-transformed red runner prey body mass (*log*(*cap time*) *∼ trial temp* + *log*(*mass*). Statistical text boxes display the OLS coefficient of determination (*R*^2^), two-tailed Student’s t-test p-values for the temperature effect, and the trial sample size (n) for each animal.

**Figure S6:**
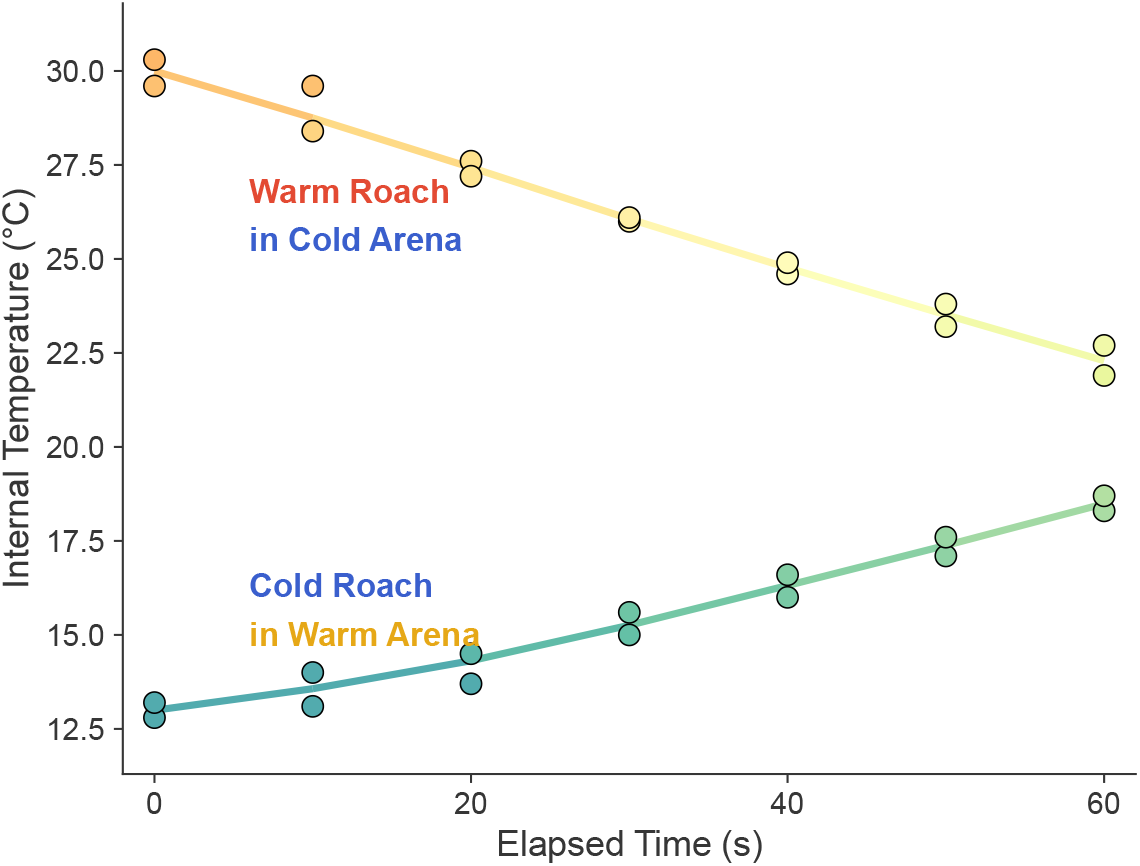
Red runner internal temperature equilibrates slowly relative to the duration of a typical hunting trial. **(A)** Internal body temperature of thermally mismatched red runners as a function of time after introduction to an arena held at the opposing temperature. Warm-acclimated red runners (30*^◦^*C) placed in a cold arena (14*^◦^*C; upper curve) cooled gradually, while cold-acclimated red runners (14*^◦^*C) placed in a warm arena (30*^◦^*C; lower curve) warmed gradually. Each point is an individual red runner (*n* = 2 per time-point); solid lines are LOESS fits. Over the first 20 s — encompassing the vast majority of capture times across conditions — internal temperature changed by only a few degrees, such that mismatched prey retained body temperatures near their acclimation state for the duration of a typical trial. This limited equilibration means that the prey behavior experienced by the mouse in mismatch trials was governed by the red runner’s acclimation temperature rather than the ambient arena temperature.

**Figure S7:**
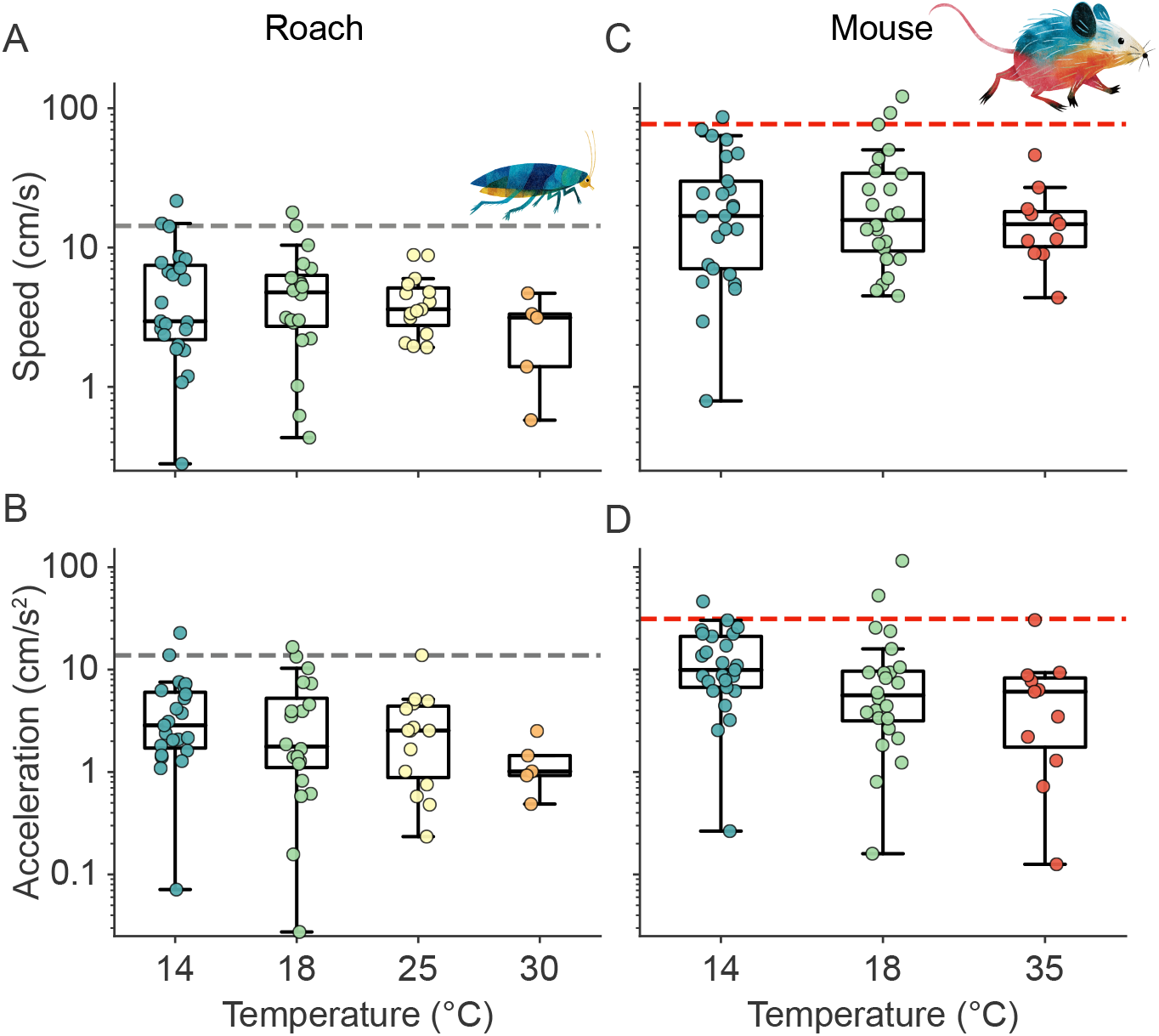
Non-locomotory movement thresholds. Speed **(A, C)** and acceleration **(B, D)** of red runners **(A, B)** and mice **(C, D)** during manually identified periods of non-locomotory movement (e.g., grooming or pose-estimation jitter), as a function of temperature. Each point is one recording (*n* = 63 red runner, *n* = 60 mouse); box plots represent medians grouped by target temperature (14, 18, 25, 30, or 35*^◦^*C) and interquartile range (IQR). Dashed lines mark the 95% percentile of non-locomotory movement for each species (red runner speed, 14.3 cm s*^−^*^1^; mouse speed, 76.9 cm s*^−^*^1^; red runner acceleration, 13.8 cm s*^−^*^2^; mouse acceleration, 31.3 cm s*^−^*^2^), used as thresholds to exclude non-locomotory movement from locomotor analyses. *y*-axes are log-scaled.

**Figure S8:**
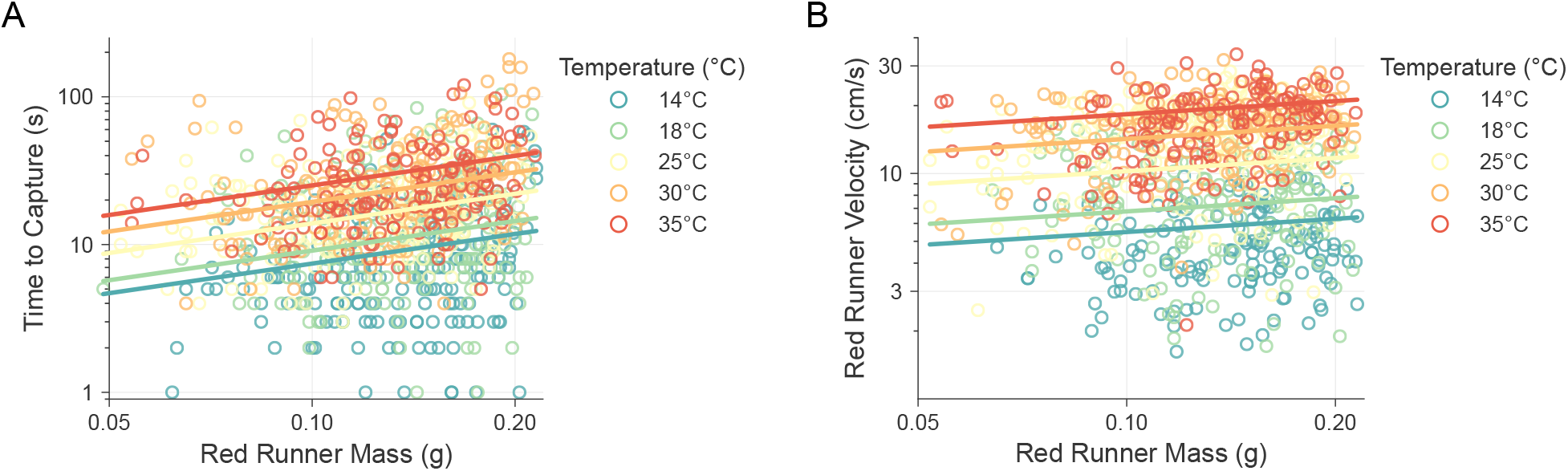
Time to capture and red runner speed as a function of red runner mass across temperatures. **(A)** Time to capture and **(B)** red runner speed during pursuit, each plotted against red runner body mass across all thermal trial hunts, colored by target temperature (*n* = 7 mice, 135.9 *±* 13.6 trials per mouse, *n* = 951 total trials). Points represent individual trials; solid lines are additive model fits evaluated at the median trial temperature for each target temperature group. Log-transformed response variables were analyzed using an LMM with mean trial temperature included as a fixed covariate and animal ID as a random intercept (*log*(*response*) *∼ log*(*mass*) + *trial temp* + (1*|animal*)). **(A)** Time to capture significantly increased with prey mass (*p <* 0.001) and trial temperature (*p <* 0.001), while accounting for significant baseline inter-individual variation across mice (random intercept, *p <* 0.001; marginal *R_m_*^2^ = 0.273, conditional *R_c_*^2^ = 0.553). **(B)** Red runner prey speed significantly increased with mass (*p <* 0.001) and trial temperature (*p <* 0.001), while accounting for baseline inter-individual variation across mice (random intercept, *p* = 0.05; marginal *R_m_*^2^ = 0.451, conditional *R_c_*^2^ = 0.468). Both axes are log-scaled in both panels.

